# Viral commitment to infection depends on host metabolism

**DOI:** 10.1101/2025.04.30.651438

**Authors:** Anastasios Marantos, Kim Sneppen, Stanley Brown, Namiko Mitarai

## Abstract

Viral infection begins with attachment to host surface structures such as receptors, pili, or porins. While prior research has focused on structural compatibility and recognition, the role of host physiology, particularly metabolic state, on viral commitment to infection remains underexplored. Here, we measured the adsorption rates (*η*) of five *Escherichia coli* phages representing various life cycles and entry pathways under controlled metabolic conditions. Four phages showed significantly reduced adsorption under energy-limited states, with weaker-binding phages being more sensitive. Using *E. coli* and its phages allowed us to institute a number of control infections that would be difficult with other organisms. Our findings support a two-step infection model where bound phages may disengage under unfavorable conditions, reducing commitment to non-productive infections. We observed a correlation between adsorption rates under energy-competent conditions and sensitivity to host metabolic state. Our results highlight host physiology as a key factor in virus–host interactions under energy-limited conditions.

## Introduction

Bacteriophages (phages) are viruses that infect bacteria, first identified by Frederick Twort and Félix d’Hérelle in the early 20th century (***Twort, 1915; d’Hérelles, 1917; Keen, 2015***). To establish infection, phages must first encounter their host (***Adam and Delbrück, 1968; Berg and Purcell, 1977***) and then their genomes must successfully enter the bacterial cell (***Tolmach, 1957; Hu et al., 2013; Bebeacua et al., 2013; Spinelli et al., 2014; Rothenberg et al., 2011; Wedd et al., 2024***). This process exploits bacterial surface structures such as outer membrane receptors, pili, and porins (***Bertozzi Silva et al., 2016***). Phage-bacteria interactions are typically studied under controlled laboratory conditions (***Madigan et al., 2018***); however, in natural environments, bacteria often experience suboptimal conditions for growth, such as nutrient limitation (e.g., nitrogen or phosphorus scarcity), fluctuating temperatures, and variable energy availability (e.g., reduced ATP production under anaerobic conditions) (***Lennon and Jones, 2011***). These conditions profoundly influence host physiology (***Lennon and Jones, 2011***) and exert strong eco-evolutionary pressures that shape phage-bacteria interactions and, by extension, microbial ecosystem dynamics (***Jones and Lennon, 2010; Shoemaker and Lennon, 2018; Igler, 2022; Măgălie et al., 2026***). A critical factor in microbial life under suboptimal conditions is the impact on host metabolic activity. While the effects of metabolism on post-infection processes, such as viral replication and lysogeny, have been extensively studied (***Kourilsky, 1973, 1974; Kourilsky and Knapp, 1974; Kourilsky and Gros, 1976; Stewart and Levin, 1984; Hadas et al., 1997; Arkin et al., 1998; You et al., 2002; Ptashne, 2004; Zeng et al., 2010; Maslov and Sneppen, 2015; Golding, 2018; Li et al., 2020; Golding et al., 2021; Geng et al., 2024; Goel et al., 2025***), its effect on viral adsorption has not received the same level of attention (***Storms and Sauvageau, 2015; Leprince and Mahillon, 2023***).

The foundations of phage adsorption studies were set by the works of Krueger, Schlesinger, Luria, Delbrück and Weidel (***Krueger and Northrop, 1930; Krueger, 1931; Delbrück, 1940; Luria and Delbrück, 1943; Weidel, 1951; Weidel and Kellenberger, 1955; Weidel, 1958***). First hypothesized by Anderson and later experimentally demonstrated for the first time by Garen and Puck for phage T1, viral entry occurs in two distinct steps: an initial, reversible attachment that does not require energy, followed by an irreversible, energy-dependent binding step (***Anderson, 1949; Garen and Puck, 1951; Stent and Wollman, 1952***). This mechanism was later corroborated and extended through the Institute Pasteur group’s seminal studies on phage *λ* and its interaction with the LamB protein receptor (***Randall-Hazelbauer and Schwartz, 1973; Schwartz and Le Minor, 1975; Schwartz, 1975; Szmelcman and Hofnung, 1975; Schwartz, 1976, 1980***), as well as Volkmar Braun et al.’s pioneering work on the FhuA protein receptor, which serves as the entry point for phages T1, T5, and *ϕ*80 (***Braun et al., 1973; Hantke and Braun, 1975; Hancock and Braun, 1976; Hantke and Braun, 1978; Kadner et al., 1980; Schöffler and Braun, 1989; Killmann and Braun, 1994; Killmann et al., 1995, 1996; Braun, 2009, 2018***). This two-step process exhibits interesting mechanistic nuances. Braun’s research established that FhuA functions as a gated channel requiring energy for phage entry (***Schöffler and Braun, 1989; Killmann and Braun, 1994; Killmann et al., 1995, 1996***). While T1 and *ϕ*80 adsorption depend on host metabolism (***Hancock and Braun, 1976; Kadner et al., 1980***), T5 can still enter energy-depleted cells (***Hantke and Braun, 1978; Kadner et al., 1980; Killmann and Braun, 1994***). This is because FhuA acts as a closed gate for the virus, requiring energy to undergo a conformational change that allows entry (***Hantke and Braun, 1978; Schöffler and Braun, 1989; Killmann et al., 1995***). In energy-depleted cells, this energy-dependent conformational change does not occur, preventing most phages from entering. However, the closure of the FhuA channel is not absolute, and T5 manages to enter (***Hantke and Braun, 1978; Kadner et al., 1980; Killmann and Braun, 1994***). Notably, we observed that for phage *λ*, adsorption is also sensitive to the metabolic state of the host (***Brown et al., 2022***) and is inhibited by the same compounds that block LamB hyperdiffusion (***Winther et al., 2009***). Moreover, a mutant phage variant (*λh*) could bypass this dependency ***Brown et al***. (***2022***), indicating that adsorption by wild-type *λ* is not solely determined by receptor presence but is also modulated by host energy availability in a non-trivial way. Furthermore, recent single-cell observations showed that phage entry probability decreases when the multiplicity of infection is high, due to changes in membrane integrity and loss of membrane potential (***Nguyen et al., 2024***). Note that these energy dependences of phage adsorption are beyond what could be explained by bacteria cell size variation due to growth conditions (***Hadas et al., 1997***).

Although these studies have independently explored the influence of host metabolism on phage adsorption, the phenomenon was rarely a central focus and often emerged as a secondary or incidental observation. This may partly explain why it has not yet received broader attention within the microbial ecology and microbiology communities (***Storms and Sauvageau, 2015; Leprince and Mahillon, 2023***), especially considering its implications for population-level virus–host interactions in environments where energy-limited conditions are the norm (***Lennon and Jones, 2011***).

Here, we aim to bring this phenomenon back into focus by providing, for the first time to our knowledge, a systematic and quantitative comparison across different phages and entry pathways to test the generality and comparative effect of host’s metabolism on viral adsorption. In this study, we investigated how the host’s metabolic state influences bacteriophage commitment to infection as indicated by the loss of the phage’s ability to infect newly introduced host bacteria. In this survey we examined a diverse set of *Escherichia coli* phages with different life cycles and entry mechanisms. By utilizing a common standardized experimental protocol, we provided a direct comparative analysis of adsorption efficiencies under identical conditions and quantified the dependence of phage adsorption rates on host metabolic state.

Our results demonstrate that metabolic-state-dependent commitment to infection is a widespread phenomenon, observed in four out of the five phages studied. This holds true across phages with diverse infection strategies, including lytic, lysogenic, and chronic phages, and entry pathways as diverse as LamB and FhuA receptors, the bacterial pilus, and the Tsx porin. To enable direct quantitative comparisons, we measured the adsorption rate constant (*η*) for each phage under standardized conditions, providing one of the few comparative accounts of *η* across multiple phages. We further quantified how host metabolism affects adsorption efficiency and found a strong correlation: phages with higher baseline *η* values under energy-rich conditions were less sensitive to metabolic inhibition.

## Results

To assess the impact of the host’s metabolic state on viral commitment to infection, we exposed each phage to bacteria under both energy-competent (in the presence of glucose (from now on denoted as Glu)) and energy-depleted conditions (in the presence of the metabolic inhibitors potassium arsenate and sodium azide (from now on denoted as As/Az)), as described in *Methods and Materials*. By quantifying the viral particles as plaque-forming units (PFU) in the post-cellular supernatant and comparing permissive hosts to resistant hosts and buffer controls under each condition, we measured the commitment to infection for each metabolic state (as shown in Tables S1 and S2).

To illustrate this effect, we define two normalized measures of the number of free viral particles after mixing with bacteria under each metabolic condition:

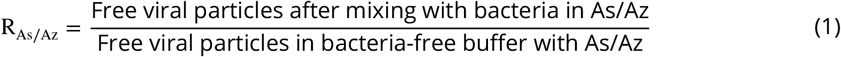

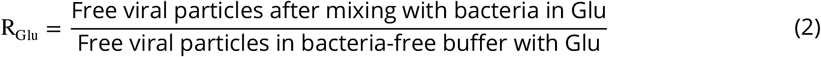

Here, R_As/Az_ represents the relative abundance of free viruses under energy-depleted conditions, and R_Glu_ represents the corresponding value under energy-competent conditions. The ratio of these two quantities captures how the host’s metabolic state influences viral commitment to infection. We define:

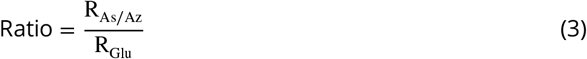

This analysis was performed for both permissive and resistant hosts, with the resistant strain serving as a control for the effects of the conditions on adsorption. The results of these experiments are visualized in Figure 1 and detailed in Tables S1, S2 and Figures S1, S2.

**Figure 1.**
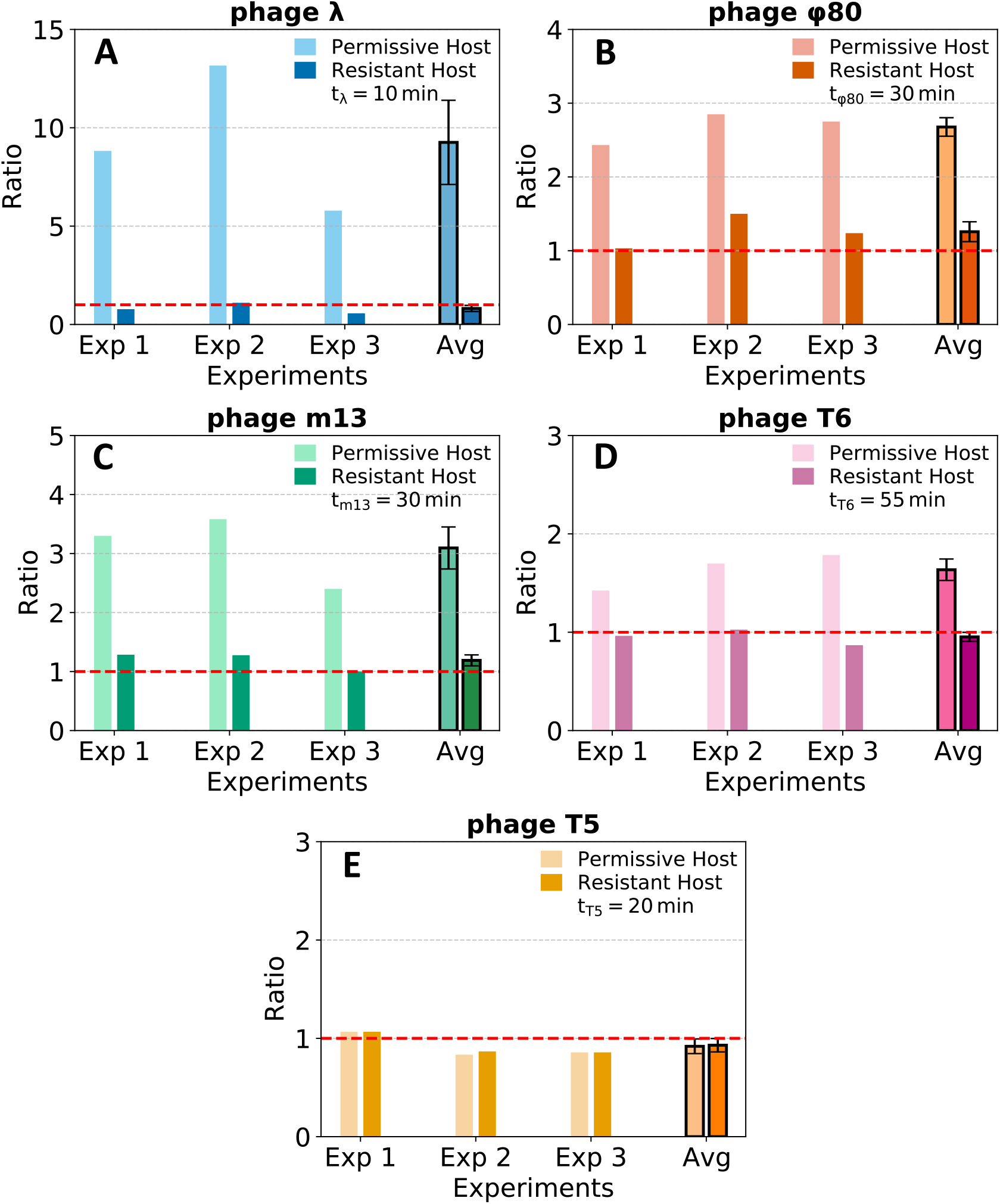
Effect of hosts’ metabolic condition on viral commitment to infection. Panels **A–E** display the results of the Ratio, defined as 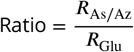, where *R*_As/Az_ =(Free viral particles after mixing with bacteria in As/Az)/(Free viral particles in bacteria-free buffer with As/Az), and *R*_Glu_ = (Free viral particles after mixing with bacteria in Glu)/ (Free viral particles in bacteria-free buffer with Glu). This Ratio captures how the host’s metabolic state affects viral commitment to infection, comparing energy-depleted (As/Az) with energy-competent (Glu) conditions across three experiments and their average. Each panel corresponds to a specific phage, as indicated above the respective panel. Lighter-colored bars represent data from permissive hosts, while darker-colored bars show results from resistant host controls for comparison. The red horizontal dashed line represents the scenario where Ratio = 1, indicating that the number of free viral particles is the same in energy-competent and energy-depleted bacteria. The incubation time for each phage-host pair is consistent across all three experiments and is displayed in the upper-right corner of each panel. The standard error of the mean (SEM) is used to estimate the variability in the averages, accounting for the random measurement errors across the three independent experiments.

If the metabolic state of the host had no effect on viral commitment, as seen in the resistant control case, the Ratio for permissive strains would be equal to 1. A Ratio greater than 1 indicates that more free viruses remain when exposed to energy-depleted hosts compared to energy-competent hosts (for example, a Ratio = 2 corresponds to twice as many free viruses), whereas a Ratio less than 1 indicates fewer free viruses under energy-depleted conditions (for example, a Ratio = 0.5 corresponds to half as many free viruses).

For the temperate lambdoid phages *λ* and *ϕ*80, the Ratio values were 9.3 and 2.7, respectively (Figures 1A–B). The chronic bacteriophage m13 showed a Ratio greater than 3 (Figure 1C), and the virulent phage T6 had a Ratio of approximately 1.6 (Figure 1D). In contrast, for phage T5, the Ratio remained close to 1 (Figure 1E), consistent with the absence of variation between metabolic conditions.

It is important to note that the ratio measure is not directly comparable between different phages. As described in the *Methods and Materials* section, the incubation time varied for each phage strain (displayed in the top right corner of each panel in Figure 1). This variation was necessary to maximize the incubation period while ensuring that manipulations were completed before the first infection cycle concluded. By allowing sufficient time for phage adsorption but preventing completion of a bacteriophage growth cycle, we ensured that only first-generation viruses were present. However, to make the effect of the host’s metabolic state comparable across different phages, we need to eliminate the influence of time dependency.

To achieve this, we employed a standard mass-action kinetics model of phage adsorption (***Krueger and Northrop, 1930; Krueger, 1931; Schlesinger, 1932; Delbrück, 1940***), where free viruses are removed from the medium through probabilistic random encounters with bacteria at a rate proportional to their concentration:

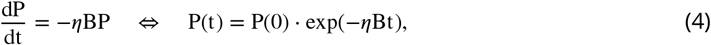

where B = (3 ± 0.5) × 10^9^ cells/mL represents the bacterial density, assumed to remain constant throughout the relevant experimental period, as bacterial growth and lysis were negligible after bacterial resuspension in buffer. Here, P(0) represents the initial viral concentration in the buffer, while P(t) corresponds to the viral concentration at a given time. Therefore, the only factor differentiating the number of “Free viral particles in Glucose” (P_GLU_(t)) and “Free viral particles in Arsenate and Azide” (P_As/Az_(t)) for each phage is the adsorption rate, which we can now determine by solving

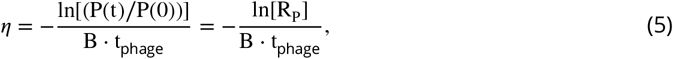

where t_phage_ denotes the incubation time allowed for each respective phage.

Crucially, this approach also allows us to compare the effect of different metabolic conditions on adsorption across different phages, as this measure is now independent of the duration of time over which we measure phage adsorption. Defining *η*^′^ as the adsorption rate in arsenate and azide, we can express the relative change in adsorption as:

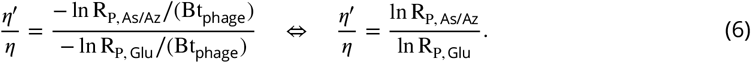

The results for each phage are summarized in Table 1, and analyzed in more detail in Table S2. The adsorption rates (*η*) were determined for each phage in glucose-grown cultures and compared with a range of adsorption rate values from the literature. While these comparisons provide useful context, caution is necessary, as differences in methodology, including experimental setup, bacterial and phage strains, and culturing conditions, can substantially influence adsorption rates. Such methodological variations have been shown to alter adsorption rates by an order of magnitude or more and occasionally even exceed a 100-fold difference (***Heller and Braun, 1979; Moldovan et al., 2007; Braun, 2009; Storms et al., 2012; Tomat et al., 2022***). For example, in the case of T6, where we observe the largest discrepancy, literature values were obtained using different bacterial strains, media composition, and a significantly lower experimental temperature (24°C compared to our 37°C).

**Table 1.** Adsorption rate in glucose (*η*) and the effect of altered metabolic conditions (growth in arsenate and azide) on it (*η*^′^/*η*) for different phages. This table presents the experimentally determined adsorption rates (*η*) for each phage when grown in glucose, along with the relative adsorption rates (*η*^′^/*η*) under arsenate and azide (As/Az) conditions, reflecting the impact of altered metabolic states. In the cases of m13 and T6, the relative adsorption rate (*η*^′^/*η*) exhibits increased variability, with SEMs exceeding the mean values. This is due to very low adsorption under As/Az conditions, where free viral particle counts approach those of the buffer control (see Figure S1 and Table S2). While this inflates relative variability, the results remain consistent with a strong reduction in adsorption efficiency. Furthermore, corresponding literature value ranges for *η* are included for comparison, with references provided. However, it is important to highlight that such comparisons are limited by differences in experimental protocols, host and virus strains, and growth conditions (e.g., temperature and media). Altering any of these factors has been reported to cause variations in adsorption rates, typically by up to an order of magnitude, and in some cases by even more than a 100-fold (***Heller and Braun, 1979; Moldovan et al., 2007; Braun, 2009; Storms et al., 2012; Tomat et al., 2022***). For example, in the case of T6, where we observe the largest discrepancy, literature values were obtained using different bacterial strains, grown in different media, at 24°C, which is 13°C lower than the 37°C temperature used in our experiments. ∗^1^:(***Hendrix and Duda, 1992; De Paepe and Taddei, 2006; Moldovan et al., 2007; Shao and Wang, 2008; Storms et al., 2012***). ∗^2^:(***Kadner et al., 1980; De Paepe and Taddei, 2006***). ∗^3^:(***Tzagoloff and Pratt, 1964; De Paepe and Taddei, 2006***). ∗^4^:(***Storms et al., 2012***). ∗^5^:(***Heller and Braun, 1979; Kadner et al., 1980; De Paepe and Taddei, 2006; Storms et al., 2012***).

| Phage | $\eta$ ($\times 10^{-11}$ mL/(CFU·min)) | $\eta'/\eta$ | Literature<br>$\eta$ ($\times 10^{-11}$ mL/(CFU·min)) | References |
| --- | --- | --- | --- | --- |
| $\lambda$ | $15.5 \pm 2.7$ | $0.53 \pm 0.03$ | [1.3, 1100] | *<sup>1</sup> |
| $\phi 80$ | $1.7 \pm 0.3$ | $0.37 \pm 0.05$ | [11, 38] | *<sup>2</sup> |
| m13 | $1.32 \pm 0.23$ | $0.06 \pm 0.08$ | [3, 9] | *<sup>3</sup> |
| T6 | $0.32 \pm 0.06$ | $0.08 \pm 0.11$ | [93, 340] | *<sup>4</sup> |
| T5 | $5.8 \pm 1.2$ | $1.03 \pm 0.16$ | [4, 2500] | *<sup>5</sup> |

For all phages tested, the relative adsorption rate (*η*^′^/*η*) under arsenate and azide (As/Az) conditions indicated a reduction in adsorption efficiency. In the cases of m13 and T6, this effect was especially pronounced, with the standard error of the mean (SEMs) for *η*^′^/*η* exceeding the mean values. This variability arises from extremely low adsorption under As/Az conditions, where the number of free viral particles in the supernatant approached those of the buffer control (see Figure S1 and Table S2). Despite this, the trend remains consistent with a substantial impairment of adsorption efficiency under metabolic stress.

Furthermore, in Figure 2, we visualize the impact of metabolic conditions on adsorption rate across different phages. In panel A, we compare the ratio *η*^′^/*η* for each phage, while in panel B, we examine the relationship between metabolic condition sensitivity (*η*^′^/*η*) and the baseline adsorption rate in glucose (*η*) of each phage. In particular, among phages that exhibit metabolic state sensitivity, we observe a significant correlation between *η*^′^/*η* and log(*η*), with a Pearson correlation coefficient of *r* = 0.86, although with *r*^2^ = 0.74 and an orthogonal distance regression fit that yields 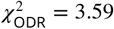.

**Figure 2.**
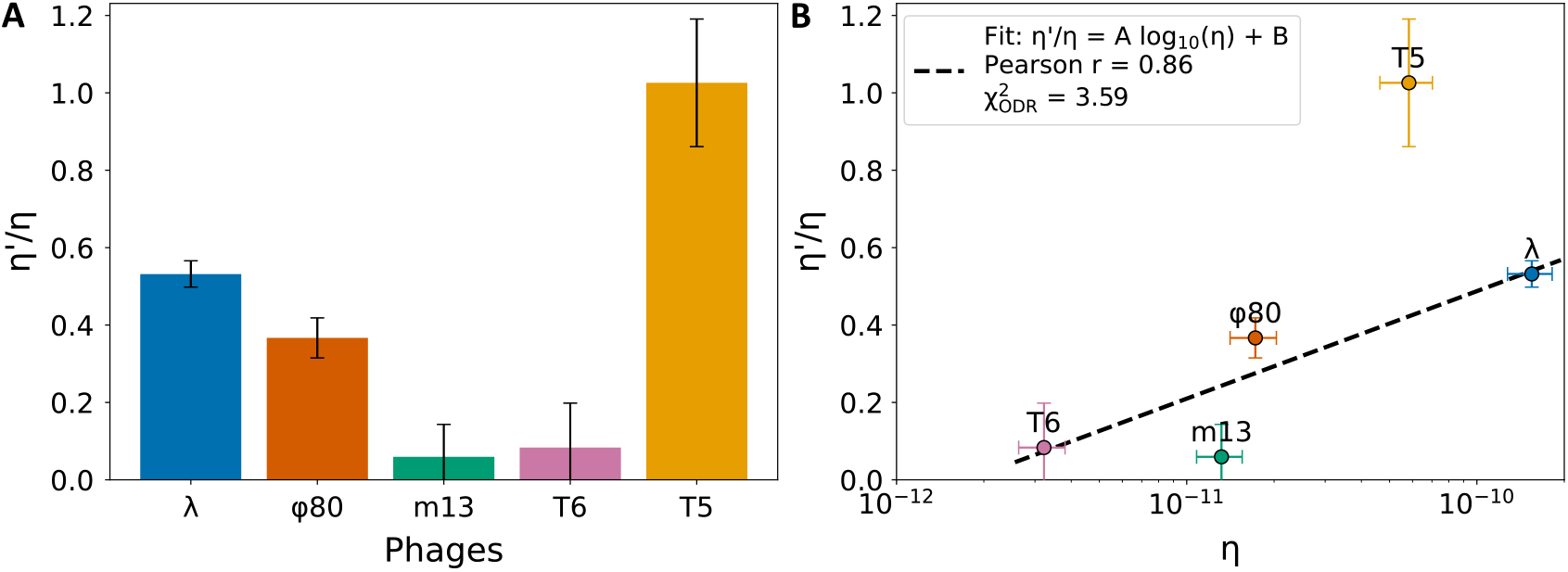
Comparing the metabolic condition effect on phage adsorption rate. The y-axis of both panels represents the relative effect of growth in arsenate and azide (low metabolic condition) on the adsorption rate, *η*^′^, compared to the adsorption rate in the high metabolic condition, *η*, for hosts grown in glucose. Panel **A** displays this effect for each phage (x-axis), while panel **B** illustrates the relationship between the effect and the adsorption rate in glucose, *η*. The dashed line in panel **B** shows an orthogonal distance regression (ODR) fit of the phages whose adsorption was affected by the host’s metabolic state (T5 was excluded because no detectable effect was observed). The relationship is modeled as *η*^′^/*η* = *A* log_10_(*η*) + *B*, with fitted parameters *A* = 0.28 ± 0.10 and *B* = 3.3 ± 1.0, where the uncertainties denote standard errors of the fit. The plotted regression line uses the full-precision parameter values, while the values reported here are rounded for clarity.

## Discussion

We sought to determine whether the efficiency of viral entry is modulated by the metabolic state of the bacterial host. In the process we explored the generality of this phenomenon in a quantitative way that would allow for comparison between different phages. To address this, we exposed a diverse set of *E. coli* phages to bacterial cells under contrasting energy conditions and quantified the proportion of phages that failed to bind irreversibly to energy-depleted hosts, while retaining the ability to infect newly introduced hosts in an energy-competent state. Specifically, we measured the concentration of free phage particles remaining in the supernatant after a controlled adsorption period, using plaque assays to estimate the number of plaque-forming units (PFUs). Because only unadsorbed phages remain in the supernatant at this stage, and since the entire assay is completed before completion of the phage growth cycle, the PFU count serves as a proxy for binding efficiency. By comparing PFU counts across energy-competent and energy-depleted conditions and normalizing them against buffer and resistant host controls, we quantified the extent to which entry depends on host metabolic activity. Using the *E. coli*–phage model was essential for implementing the control infections, as its genetic and physiological tractability enables a level of experimental precision that is difficult to achieve in other systems.

This behavior is further illustrated in the schematics provided in Figure 3, which summarizes the observed phenomenon. At the individual level, a phage moves through the environment in a random walk until it encounters a host. Our results reveal that if the host is in a high metabolic state (energy-competent), viral commitment occurs. In contrast, when the bacterium is in a low metabolic state (energy-depleted), the phage does not bind irreversibly to the host. At the population level, this metabolic-state-dependent infection results in a higher proportion of free viral particles when bacteria are grown in conditions that induce low metabolic activity (e.g., arsenate or azide) compared to bacteria grown in glucose, which supports higher metabolic activity. Thus, the fraction of phages committing to infection is greater when encountering energy-competent hosts than when encountering energy-depleted hosts.

**Figure 3.**
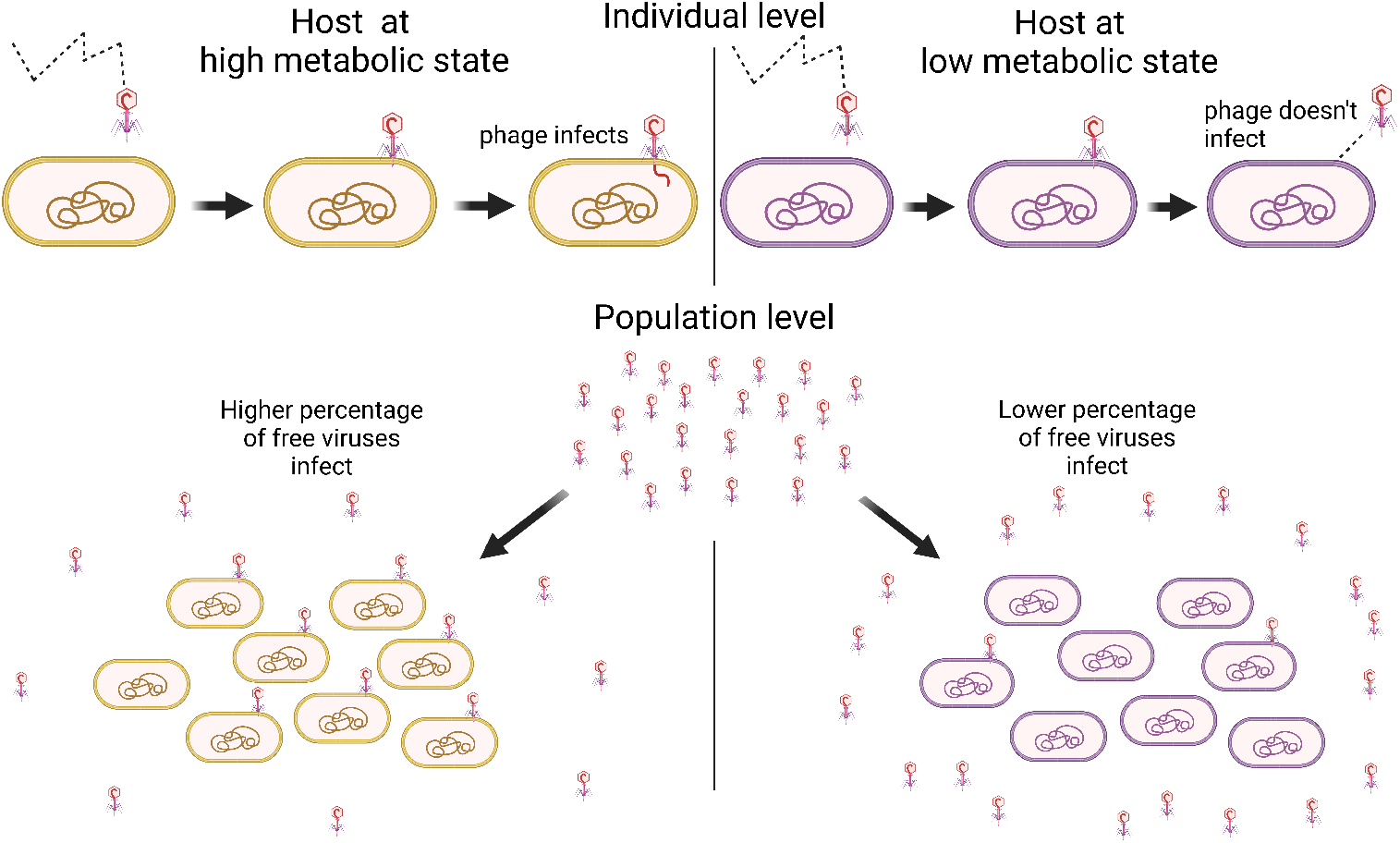
Schematic of the Phenomenon. This illustration compares the ability of viral particles to enter upon encountering energy-competent bacteria (yellow, left) versus energy-depleted bacteria (purple, right). The upper part depicts a sequence of events, following the arrows from left to right, showing phage behavior when encountering a high-metabolic-state (energy-competent) host versus a low-metabolic-state (energy-depleted) host. At the population level, a greater percentage of free viral particles in the buffer will commit to infecting a community of high-metabolic-state hosts (grown in glucose) compared to those at a low metabolic state (grown in arsenate and azide).(Created in BioRender. Marantos, A. (2025) https://BioRender.com/3ymwbkk).

Further, by estimating the phage-specific adsorption rate constants (*η*), we corrected for differences in incubation time, enabling time-independent comparison across phage types. Our approach employs a simple, reproducible, and quantifiable framework in which all phages are studied in the same laboratory under consistent protocols, yielding directly comparable results across different viruses. The phenomenon exhibits remarkable generality, manifested across a variety of viral types, from lysogenic lambdoid phages (*λ* and *ϕ*80), to chronic phages (m13), and virulent phages (T6). Furthermore, these phages utilize distinct entry pathways to exploit the host. For instance, phage *λ* relies on the maltose uptake pathway through LamB (***Randall-Hazelbauer and Schwartz, 1973***), *ϕ*80 uses the iron uptake pathway through FhuA and TonB (***Hantke and Braun, 1975; Hancock and Braun, 1976; Kadner et al., 1980; Ferguson et al., 1998***), T6 exploits the Tsx porin (***Nakae, 1976; Hantke, 1976; Manning and Reeves, 1978; Bremer et al., 1988, 1990***), and m13 utilizes the pilus (***Tzagoloff and Pratt, 1964; Madigan et al., 2018***).

Interestingly, despite this generality, the phenomenon does not apply to all phages. T5 does not exhibit dependence on the metabolic state of the host for entry, even though it also utilizes FhuA as its entry receptor (***Braun et al., 1973; Luckey et al., 1975; Hantke and Braun, 1975, 1978; Ferguson et al., 1998***), similar to *ϕ*80 (***Kadner et al., 1980; Poon and Dhillon, 1987; Killmann and Braun, 1994; Killmann et al., 1995***). This is consistent with the findings of Braun et al., who showed that the differential behavior of *ϕ*80 and T5 can be attributed to their reliance (or lack thereof) on the FhuA–TonB energy transduction system (***Hancock and Braun, 1976; Hantke and Braun, 1978; Kadner et al., 1980; Killmann and Braun, 1994; Killmann et al., 1995***). TonB is part of the host’s machinery for the uptake of ferrichromes and other compounds (***Postle, 1993; Postle and Kadner, 2003; Postle and Larsen, 2007***) and is required to energize FhuA during phage entry in the case of *ϕ*80 and T1 (***Hancock and Braun, 1976; Kadner et al., 1980; Killmann and Braun, 1994; Killmann et al., 1995***). However, under the experimental conditions used here, T5 circumvents this requirement (***Hantke and Braun, 1978***). This suggests that in energy-depleted cells, where TonB function is compromised, phages dependent on TonB cannot trigger the conformational changes required to transition from reversible binding to the irreversible, energy-dependent state and proceed with DNA injection (***Killmann and Braun, 1994; Killmann et al., 1995***). In contrast, phages such as T5, which bypass this gate, can infect unimpeded. It remains unclear whether viruses themselves possess metabolic state-sensing capabilities, but Braun et al.’s findings suggest that this modulation arises primarily from host physiology (***Braun, 2009, 2018***).

In contrast, our findings suggest that phage *λ* is capable of sensing the metabolic state of its host (***Brown et al., 2022***). These results are particularly relevant to the current study, as we used the same protocol and observed consistent behavior from phage *λ*. Importantly, our previous study (***Brown et al., 2022***), along with the experiments described in this work and the cited studies by the Institut Pasteur group and Moldovan et al. (***Randall-Hazelbauer and Schwartz, 1973; Schwartz and Le Minor, 1975; Schwartz, 1975; Szmelcman and Hofnung, 1975; Schwartz, 1976, 1980; Moldovan et al., 2007***), used *λ*wt (also known as *λ*PaPa (***Hendrix and Duda, 1992***)). This is not the original *λ* isolated in 1951 (***Lederberg, 1951***) (also known as Ur-*λ* (***Hendrix and Duda, 1992***)), but a laboratory derivative that has lost its accessory tail fibers, resulting in a lower adsorption rate and, consequently, the formation of larger plaques (***Hendrix and Duda, 1992***). In Table 1, the upper range of the *η* literature values for *λ* is set by Ur-*λ* ([600, 1100] ×10^−11^ mL min^−1^ (***Hendrix and Duda, 1992; Storms et al., 2012***)), while the lower range corresponds to the *λ*PaPa strain ([1.3, 990] ×10^−11^ mL min^−1^ (***De Paepe and Taddei, 2006; Moldovan et al., 2007; Shao and Wang, 2008***)). Phage *λ*, which requires the LamB protein for recognition and entry, was found to rely on the metabolic state of the host. We interpreted this behavior through *λ*’s reversible binding mechanism (***Schwartz, 1975***) and the role of LamB hyperdiffusion (i.e., the enhanced lateral motion of LamB proteins within the outer membrane) under energy-sufficient conditions (***Oddershede et al., 2002; Winther et al., 2009***). While wild-type *λ* relies on conditions allowing hyperdiffusion for commitment to infection, we observed that a mutant variant, *λh*, which does not depend on them, was able to infect even energy-depleted hosts (***Brown et al., 2022***). Furthermore, our data indicate that this phenomenon is not due to direct effects of arsenate or azide on phage physiology or to changes in host receptor abundance, as the mutant *λh* remained unaffected by these treatments and could infect the host as if the conditions were glucose-rich (***Brown et al., 2022***).

Because arsenate and azide are well-established inhibitors of cellular energy metabolism, they serve as effective tools for studying this phenomenon. The fact that a mutation in the tail fiber of *λh* disables the phage’s ability to differentiate hosts based on metabolic state suggests that this sensitivity may be an evolved function. Notably, wild-type *λ* is inactivated by *E. coli* K-12 extracts only when solvents are added, whereas *Shigella* extracts inactivate *λ* without this requirement (***Randall-Hazelbauer and Schwartz, 1973; Schwartz, 1975; Schwartz and Le Minor, 1975***). This suggests that *E. coli* LamB requires a specific state for irreversible binding, a conditionality absent in *Shigella* LamB, indicating that the capacity for metabolic-state sensing may depend on receptor-specific properties. Such mechanistic insights are currently feasible only for phages like *λ*, whose infection cycles are relatively well understood. Nevertheless, *λ* serves as a valuable model for understanding a broader and generalizable phenomenon.

In addition to establishing the generality of this phenomenon, our study contributes one of the few comparative accounts of directly measured adsorption rate constants (*η*) under standardized conditions. These rates varied considerably across phages, and when comparing *η* under high and low metabolic states, we observed substantial variation in sensitivity. Notably, the degree of adsorption reduction under metabolic inhibition—captured by the ratio *η*^′^/*η*—was strongly correlated with the baseline adsorption rate in glucose-rich conditions. Phages with higher baseline *η* values (e.g., *λ*) were less sensitive to metabolic suppression, whereas those with lower adsorption rates (e.g., m13) were more affected. This relationship implies that phages capable of rapid and strong binding are more committed to infection, regardless of host state, while others may “sample” the host and disengage if conditions are suboptimal.

A two-step infection model with a discrimination mechanism offers a useful lens to understand the underlying process (see Figure 4 and references (***Stent and Wollman, 1952; Schwartz, 1976; Moldovan et al., 2007***)). In this framework, free phages *P* first bind reversibly to host cells *B* with a rate constant *k*, forming a transient bound complex [*P B*]. From this state, the phage either proceeds to commit irreversibly to the infection at rate *k*_com_, or unbinds and returns to the free state with rate *k*_off_.

**Figure 4.**
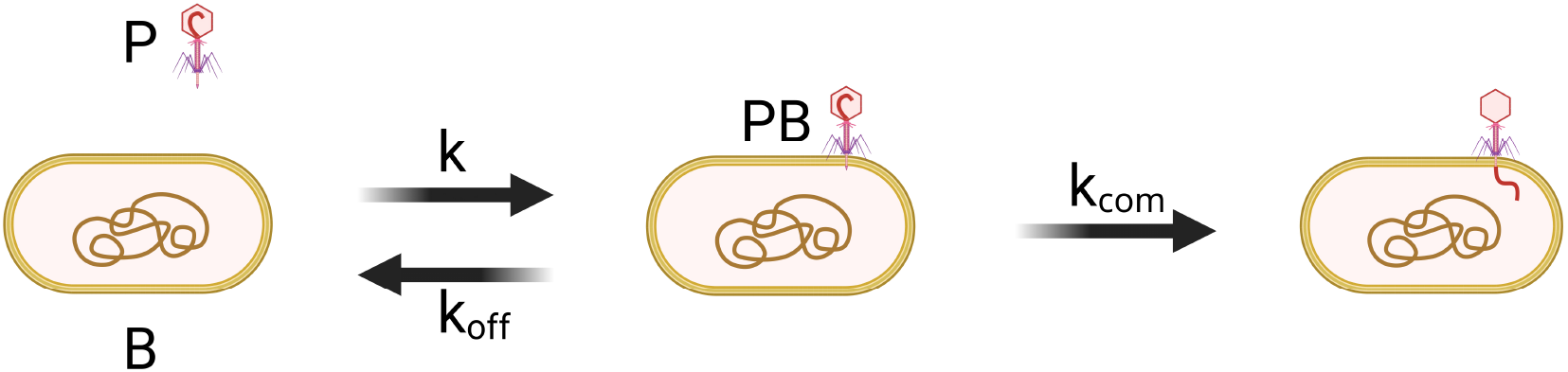
2-step infection dynamics with a discrimination mechanism. After binding to a receptor at a rate *k*, the phage can either irreversibly commit to the adsorption at a rate *k*_com_, or can leave again with a rate *k*_off_ as introduced in (***Stent and Wollman, 1952; Schwartz, 1976; Moldovan et al., 2007***). This slows down the commitment process by factor 1/(1 + *k*_off_/*k*_com_), but opens for the discrimination between the active and inactive hosts by having different values of *k*_off_/*k*_com_. (Created in BioRender. Marantos, A. (2025) https://BioRender.com/1mhwucl).

Assuming that the bound state reaches a quasi-steady-state 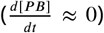, the rate of formation of the bound complex is balanced by the combined rates of dissociation and irreversible commitment:

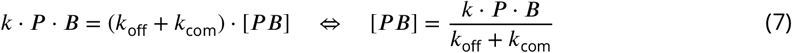

The rate of successful infection is determined by the number of complexes that proceed to irreversible DNA injection:

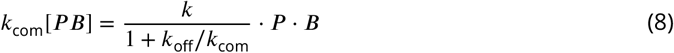

This represents the rate at which free phages are irreversibly removed from the system. Considering a small time interval Δ*t*, the fraction of phages lost is approximately:

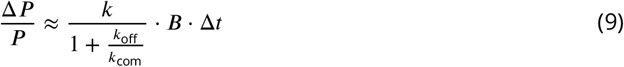

Connecting this to our mass-action model kinetics of phage adsorption, we have:

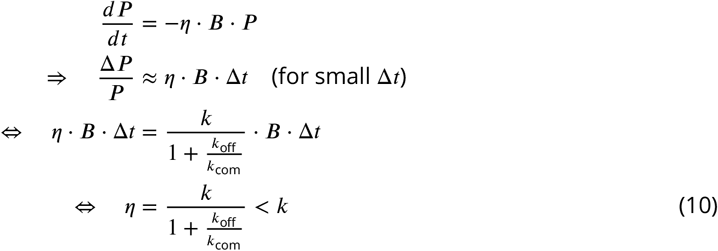

This shows that the effective adsorption rate *η* is always lower than the initial encounter rate *k* due to the possibility of unbinding before commitment to infection. However, if the phage can sense the bacteria’s physiological state and alter the value of *k*_off_/*k*_com_ accordingly, it provides an opportunity to avoid commiting to the inactive, possibly dead, host cell. Note that value of *k* is independent of the host physiological state, since the phage need to be in contact with the receptor to sense the host’s state.

Suppose a phage uses the rates *k*_off_ and *k*_com_ when encountering a metabolically active host, while it uses the rates 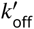 and 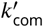 for an inactive host. Then, according to eq. (10), the resulting ratio of the adsorption rate is

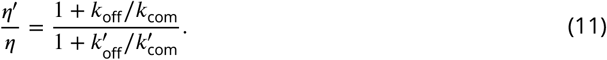

a ratio investigated in Fig. 2B. This means that the smaller the active ratio *k*_off_/*k*_com_, the bigger the inactive ratio needs to be, in order to discriminate. This illustrate an interesting trade-off: To avoid reducing *η* for a metabolically active host it is ideal to make *k*_off_/*k*_com_ as small as possible. However, a small *k*_off_/*k*_com_ requires a big 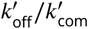 to reduce *η*^′^ ≪ *η*. This trade-off between big *η* and small *η*^′^/*η* is observed in Fig. 2B.

The ratio can be varied in two ways, corresponding to the kinetic discrimination and the energetic discrimination (***Sartori and Pigolotti, 2013***). In our framework, the kinetic discrimination corresponds to having the same off rate 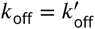 (i.e. the equilibrium binding energy to the receptor is the same) but having different commitment rate 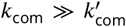. This dissipates extra energy to drive the non-equilibrium process differently. In contrast, the energetic discrimination requires the different equilibrium binding energy between the active and inactive host to ensure 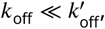, but the commitment rate are the same 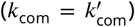. In order to the energetic discrimination to work, the commitment rate needs to be slow enough so that unbinding can actually happen, hence it costs the reaction speed.

These results support the idea that host physiology acts not only as a barrier but also as a selective filter that influences phage infection strategies. This aligns with our findings and suggests that phages may exploit host energy levels as an internal “checkpoint”, committing to infection only when the host is in a metabolically favorable state. These infection “checkpoints” may represent an elegant evolutionary strategy to minimize non-productive infection. However, such a check-point mechanism entails a fitness cost as some phages may leave viable hosts, but likely helps avoid wasteful infection attempts in poor environments. Conversely, phages like T5, which readily infect metabolically inactive cells, may be at a net disadvantage in niches containing a mixture of energy-competent and energy-depleted hosts. This is consistent with previous findings that T5 can replicate in starved bacterial cells (***Vidakovic et al., 2018***).

Together, our findings support the view that phage-host interactions are not solely determined by receptor compatibility but are dynamically modulated by host physiology. Host energy availability emerges as a key determinant of infection success at the earliest stages of the viral life cycle. This insight may help explain variation in phage efficacy in natural and clinical settings, where bacterial populations exhibit substantial physiological heterogeneity (***Pearl et al., 2008; Himeoka and Mitarai, 2020; Kolter et al., 2022; Mitarai et al., 2023***). In conclusion, we provide direct evidence that viral commitment to infection is shaped by the metabolic condition of the bacterial host. This effect, while widespread, varies in magnitude and mechanism. By combining a unified experimental framework with adsorption kinetics modeling, we reveal a physiologically gated layer of phage infection that likely contributes to both ecological dynamics and evolutionary strategies. Our work underscores the importance of incorporating host physiology into models of viral infection, especially in contexts where metabolic diversity is the norm.

## Methods and Materials

This experiment examined phage adsorption to *Escherichia coli* cells under two contrasting metabolic states: a state of high metabolic activity due to the presence of glucose, and a state of low metabolic activity due to the presence of potassium arsenate and sodium azide. The following protocol was adapted from the experimental methodology of the study by Brown et al. (***Brown et al., 2022***) (see Figure 5 and Supplementary Information for greater detail).

**Figure 5.**
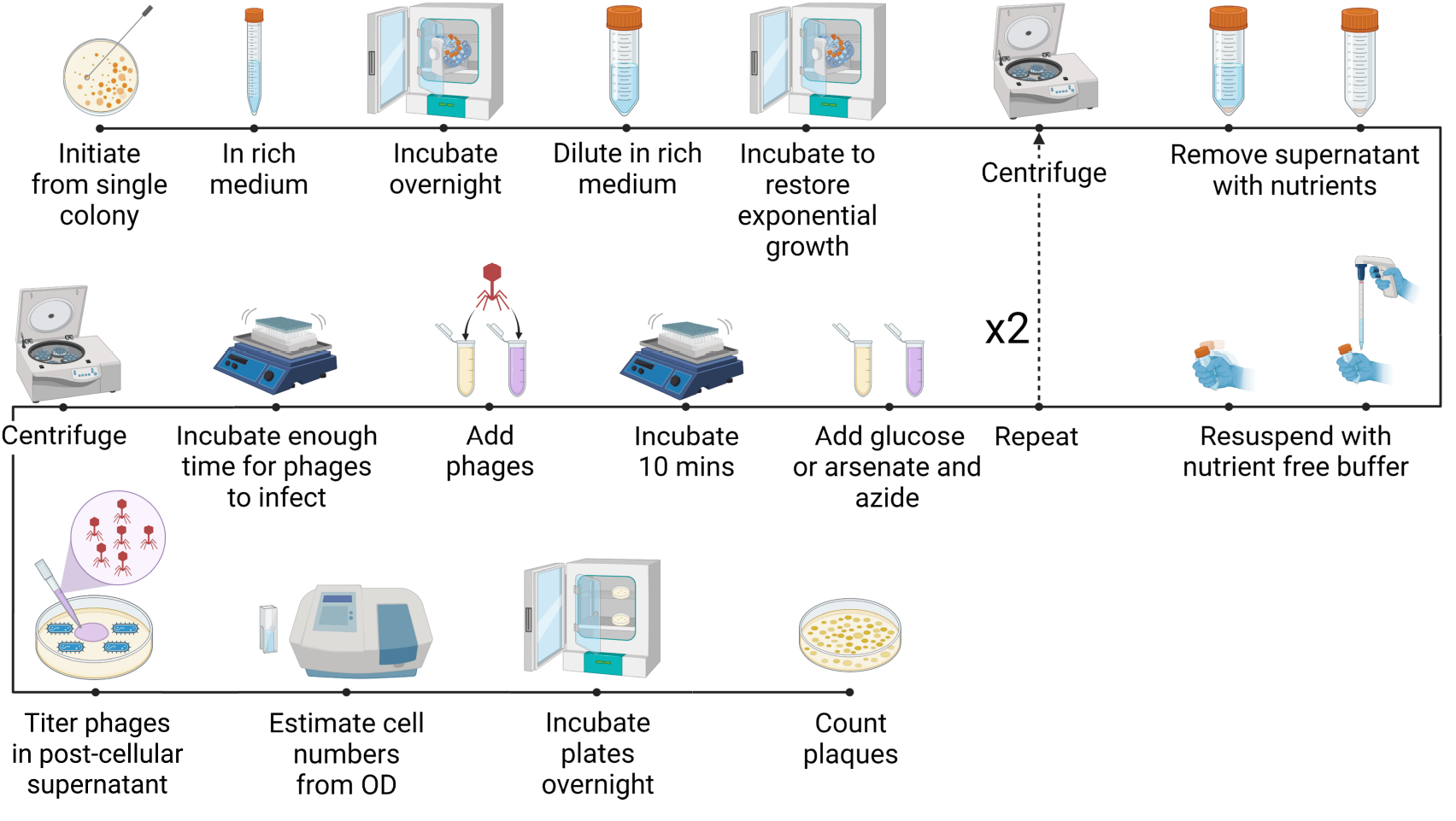
The experimental protocol. Each dot represents a step in the experimental protocol. The accompanying text below and the image above provide a detailed description of the process. The sequence of steps follows the solid line from the upper left (start) to the lower right (end), indicating the correct order. The dotted line with an arrow and the “x2” marker denotes a looped process in the protocol. (Created in BioRender. Marantos, A. (2025) https://BioRender.com/2u4x65f.)

Each bacterial culture was initiated from an independent single colony, and each phage lysate was prepared from an independent plaque. Following overnight growth, the bacterial culture was diluted 20-fold in rich medium and incubated for 2 hours to restore exponential growth. Subsequently, nutrients were removed from the bacteria through repeated centrifugation and resuspension in a nutrient-free buffer.

After washing, the samples of the washed bacteria were subjected to favorable or unfavorable growth conditions by adding glucose and potassium phosphate or potassium arsenate and sodium azide, respectively, and incubating at 37 °C for 10 minutes. This duration was chosen based on prior work showing that arsenate–azide rapidly inhibits cellular energy metabolism (***Winther et al., 2009***) and that the ATP pool of log-phase *E. coli* turns over several times per second (***Holms et al., 1972***). Post incubation, phages were introduced at a low multiplicity of infection (MOI), on the order of 0.001, to reduce the likelihood of coinfections and incubation continued. We allowed sufficient time for phage infection but not for completion of a phage growth cycle, to ensure that only first-generation viruses were present. As a result, the incubation time varied for each strain, as our goal was to maximize the incubation period while being limited by the need to complete manipulations before the first infection cycle was completed. The free phages were then separated from the bacteria by centrifugation. A portion of the supernatant was subsequently diluted 10-fold with buffer, supplemented with potassium phosphate to neutralize any residual arsenate toxicity. Phage titers were assessed by serial dilutions in buffer and spotted on a lawn of a susceptible bacterial strain. The concentration of washed bacteria was estimated by turbidity at 600nm. The turbidity was calibrated with a Petroff-Hausser chamber. The proportion of unbound phage was calculated by comparing the concentration as PFU/ml in the supernatant derived from bacteria-containing samples with that of bacteria-free samples.

To ensure robust control conditions, parallel experiments were conducted under various configurations. These included: (i) samples containing bacteria resistant to phages, (ii) samples containing only phages, with no bacteria present, (iii) samples containing only bacteria, with no phages, and (iv) samples containing neither phages nor bacteria, with only buffer. These controls allowed us to account for background effects and isolate the specific interactions between phages and sensitive bacterial populations. To accurately quantify viral binding to host bacteria, we used isogenic pairs for our permissive-resistant host controls, where resistance was conferred by phage-receptor loss.

## Data and code availability

The experimental data supporting the findings of this study are provided in the Supplementary Material. The code used for data analysis is available on GitHub at https://github.com/TassosMar/Viral-commitment-to-infection-depends-on-host-metabolism.

## Author contributions statement

A.M. and S.B. conceived the study. Experiments were conducted by A.M. with assistance from S.B. and N.M. Data analysis was performed by A.M. under the guidance of N.M., K.S., and S.B. Modeling was conceived by N.M. and K.S. The project was supervised by K.S., S.B., and N.M. All authors contributed to writing, reviewing, and editing the manuscript. Funding was acquired by N.M. and K.S.

## Acknowledgments

This research was funded by the Novo Nordisk Foundation (NNF21OC0068775) and the Danish National Research Foundation (grant number DNRF170). We thank Sine Lo Svenningsen and Adrien Sarlet for providing strains and Mireia Cordero for guidance in laboratory techniques.

