## Supplementary material for "Viral commitment to infection depends on host metabolism"

**Abstract** Viral infection begins with attachment to host surface structures such as receptors, pili, or porins. While prior research has focused on structural compatibility and recognition, the role of host physiology, particularly metabolic state, on viral commitment to infection remains underexplored. Here, we measured the adsorption rates ($\eta$) of five *Escherichia coli* phages representing various life cycles and entry pathways under controlled metabolic conditions. Four phages showed significantly reduced adsorption under energy-limited states, with weaker-binding phages being more sensitive. Using *E. coli* and its phages allowed us to institute a number of control infections that would be difficult with other organisms. Our findings support a two-step infection model where bound phages may disengage under unfavorable conditions, reducing commitment to non-productive infections. We observed a correlation between adsorption rates under energy-competent conditions and sensitivity to host metabolic state. Our results highlight host physiology as a key factor in virus–host interactions under energy-limited conditions.

### Experimental data

| $\lambda$ | Experiment 1 | | | Experiment 2 | | | Experiment 3 | | | All Experiments Averages | | | $\times 10^3 \frac{PFU}{mL}$ |
|---|---|---|---|---|---|---|---|---|---|---|---|---|---|
| | Bacteria | | | Bacteria | | | Bacteria | | | Bacteria | | | |
| | Permissive | Resistant | Buffer | Permissive | Resistant | Buffer | Permissive | Resistant | Buffer | Permissive | Resistant | Buffer | |
| | S3207 | S3222 | TMG | S3207 | S3222 | TMG | S3207 | S3222 | TMG | S3207 | S3222 | TMG | |
| Virus $\lambda$ | 11 | 1500 | 1100 | 9 | 1000 | 1100 | 10 | 800 | 900 | 10.0 ± 0.6 | 1100 ± 208 | 1033 ± 67 | GLU |
| Buffer TMG | 0 | 0 | 0 | 0 | 0 | 0 | 0 | 0 | 0 | 0 | 0 | 0 | |
| Virus $\lambda$ | 150 | 1800 | 1700 | 140 | 1300 | 1300 | 90 | 700 | 1400 | 127 ± 19 | 1267 ± 318 | 1467 ± 120 | As/Az |
| Buffer TMG | 0 | 0 | 0 | 0 | 0 | 0 | 0 | 0 | 0 | 0 | 0 | 0 | |
| Ratio to Buffer | $R_P$ | $R_R$ | $-\ln R_P$ | $R_P$ | $R_R$ | $-\ln R_P$ | $R_P$ | $R_R$ | $-\ln R_P$ | $R_P$ | $R_R$ | $-\ln R_P$ | |
| | 0.010 | 1.364 | 4.605 | 0.008 | 0.909 | 4.806 | 0.011 | 0.889 | 4.500 | 0.0098 ± 0.0009 | 1.05 ± 0.15 | 4.64 ± 0.09 | GLU |
| | 0.088 | 1.059 | 2.428 | 0.108 | 1.000 | 2.228 | 0.064 | 0.500 | 2.744 | 0.087 ± 0.013 | 0.85 ± 0.18 | 2.47 ± 0.15 | As/Az |
| Ratio | 8.82 | 0.78 | | 13.16 | 1.10 | | 5.79 | 0.56 | | **9.3 ± 2.1** | **0.81 ± 0.16** | | |

| $\phi 80$ | Experiment 1 | | | Experiment 2 | | | Experiment 3 | | | All Experiments Averages | | | $\times 10^3 \frac{PFU}{mL}$ |
|---|---|---|---|---|---|---|---|---|---|---|---|---|---|
| | Bacteria | | | Bacteria | | | Bacteria | | | Bacteria | | | |
| | Permissive | Resistant | Buffer | Permissive | Resistant | Buffer | Permissive | Resistant | Buffer | Permissive | Resistant | Buffer | |
| | MC4100 | S2153 | TMG | MC4100 | S2153 | TMG | MC4100 | S2153 | TMG | MC4100 | S2153 | TMG | |
| Virus $\phi 80$ | 3400 | 16000 | 13000 | 2000 | 10000 | 10000 | 2000 | 8000 | 11000 | 2467 ± 467 | 11333 ± 2404 | 11333 ± 882 | GLU |
| Buffer TMG | 0 | 0 | 0 | 0 | 0 | 0 | 0 | 0 | 0 | 0 | 0 | 0 | |
| Virus $\phi 80$ | 7000 | 14000 | 11000 | 5700 | 15000 | 10000 | 5000 | 9000 | 10000 | 5900 ± 586 | 12667 ± 1856 | 10333 ± 333 | As/Az |
| Buffer TMG | 0 | 0 | 0 | 0 | 0 | 0 | 0 | 0 | 0 | 0 | 0 | 0 | |
| Ratio to Buffer | $R_P$ | $R_R$ | $-\ln R_P$ | $R_P$ | $R_R$ | $-\ln R_P$ | $R_P$ | $R_R$ | $-\ln R_P$ | $R_P$ | $R_R$ | $-\ln R_P$ | |
| | 0.262 | 1.231 | 1.341 | 0.200 | 1.000 | 1.609 | 0.182 | 0.727 | 1.705 | 0.214 ± 0.024 | 0.99 ± 0.15 | 1.55 ± 0.11 | GLU |
| | 0.636 | 1.273 | 0.452 | 0.570 | 1.500 | 0.562 | 0.500 | 0.900 | 0.693 | 0.57 ± 0.04 | 1.22 ± 0.17 | 0.57 ± 0.07 | As/Az |
| Ratio | 2.43 | 1.03 | | 2.85 | 1.50 | | 2.75 | 1.24 | | **2.68 ± 0.13** | **1.26 ± 0.14** | | |

| m13 | Experiment 1 | | | Experiment 2 | | | Experiment 3 | | | All Experiments Averages | | | $\times 10^3 \frac{PFU}{mL}$ |
|---|---|---|---|---|---|---|---|---|---|---|---|---|---|
| | Bacteria | | | Bacteria | | | Bacteria | | | Bacteria | | | |
| | Permissive | Resistant | Buffer | Permissive | Resistant | Buffer | Permissive | Resistant | Buffer | Permissive | Resistant | Buffer | |
| | S3190 | S2153 | TMG | S3190 | S2153 | TMG | S3190 | S2153 | TMG | S3190 | S2153 | TMG | |
| Virus m13 | 10000 | 28000 | 30000 | 10000 | 28000 | 38000 | 13000 | 35000 | 40000 | 11000 ± 1000 | 30333 ± 2333 | 36000 ± 3055 | GLU |
| Buffer TMG | 0 | 0 | 0 | 0 | 0 | 0 | 0 | 0 | 0 | 0 | 0 | 0 | |
| Virus m13 | 33000 | 36000 | 30000 | 33000 | 33000 | 35000 | 32000 | 36000 | 41000 | 32667 ± 333 | 35000 ± 1000 | 35333 ± 3180 | As/Az |
| Buffer TMG | 0 | 0 | 0 | 0 | 0 | 0 | 0 | 0 | 0 | 0 | 0 | 0 | |
| Ratio to Buffer | $R_P$ | $R_R$ | $-\ln R_P$ | $R_P$ | $R_R$ | | $R_P$ | $R_R$ | $-\ln R_P$ | $R_P$ | $R_R$ | $-\ln R_P$ | |
| | 0.333 | 0.933 | 1.099 | 0.263 | 0.737 | 1.335 | 0.325 | 0.875 | 1.124 | 0.307 ± 0.022 | 0.85 ± 0.06 | 1.19 ± 0.08 | GLU |
| | 1.100 | 1.200 | -0.095 | 0.943 | 0.943 | 0.059 | 0.780 | 0.878 | 0.248 | 0.94 ± 0.09 | 1.01 ± 0.10 | 0.07 ± 0.10 | As/Az |
| Ratio | 3.30 | 1.29 | | 3.58 | 1.28 | | 2.40 | 1.00 | | **3.1 ± 0.4** | **1.19 ± 0.09** | | |

| T6 | Experiment 1 | | | Experiment 2 | | | Experiment 3 | | | All Experiments Averages | | | $\times 10^3 \frac{PFU}{mL}$ |
|---|---|---|---|---|---|---|---|---|---|---|---|---|---|
| | Bacteria | | | Bacteria | | | Bacteria | | | Bacteria | | | |
| | Permissive | Resistant | Buffer | Permissive | Resistant | Buffer | Permissive | Resistant | Buffer | Permissive | Resistant | Buffer | |
| | Δxyl | tsx | TMG | Δxyl | tsx | TMG | Δxyl | tsx | TMG | Δxyl | tsx | TMG | |
| Virus T6 | 27000 | 37000 | 45000 | 31000 | 41000 | 50000 | 18000 | 36000 | 33000 | 25333 ± 3844 | 38000 ± 1528 | 42667 ± 5044 | GLU |
| Buffer TMG | 0 | 0 | 0 | 0 | 0 | 0 | 0 | 0 | 0 | 0 | 0 | 0 | |
| Virus T6 | 41000 | 38000 | 48000 | 40000 | 32000 | 38000 | 37000 | 36000 | 38000 | 39333 ± 1202 | 35333 ± 1764 | 41333 ± 3333 | As/Az |
| Buffer TMG | 0 | 0 | 0 | 0 | 0 | 0 | 0 | 0 | 0 | 0 | 0 | 0 | |
| Ratio to Buffer | $R_P$ | $R_R$ | $-\ln R_P$ | $R_P$ | $R_R$ | $-\ln R_P$ | $R_P$ | $R_R$ | $-\ln R_P$ | $R_P$ | $R_R$ | $-\ln R_P$ | |
| | 0.600 | 0.822 | 0.511 | 0.620 | 0.820 | 0.478 | 0.545 | 1.091 | 0.606 | 0.588 ± 0.022 | 0.91 ± 0.09 | 0.53 ± 0.04 | GLU |
| | 0.854 | 0.792 | 0.158 | 1.053 | 0.842 | -0.051 | 0.974 | 0.947 | 0.027 | 0.96 ± 0.06 | 0.86 ± 0.05 | 0.04 ± 0.06 | As/Az |
| Ratio | 1.42 | 0.96 | | 1.70 | 1.03 | | 1.79 | 0.87 | | **1.64 ± 0.11** | **0.95 ± 0.05** | | |

| T5 | Experiment 1 | | | Experiment 2 | | | Experiment 3 | | | All Experiments Averages | | | $\times 10^3 \frac{PFU}{mL}$ |
|---|---|---|---|---|---|---|---|---|---|---|---|---|---|
| | Bacteria | | | Bacteria | | | Bacteria | | | Bacteria | | | |
| | Permissive | Resistant | Buffer | Permissive | Resistant | Buffer | Permissive | Resistant | Buffer | Permissive | Resistant | Buffer | |
| | MC4100 | S2153 | TMG | MC4100 | S2153 | TMG | MC4100 | S2153 | TMG | MC4100 | S2153 | TMG | |
| Virus T5 | 400 | 17000 | 16000 | 800 | 10000 | 12000 | 200 | 4000 | 12000 | 467 ± 176 | 10333 ± 3756 | 13333 ± 1333 | GLU |
| Buffer TMG | 0 | 0 | 0 | 0 | 0 | 0 | 0 | 0 | 0 | 0 | 0 | 0 | |
| Virus T5 | 400 | 17000 | 15000 | 1000 | 13000 | 18000 | 200 | 4000 | 14000 | 533 ± 240 | 11333 ± 3844 | 15667 ± 1202 | As/Az |
| Buffer TMG | 0 | 0 | 0 | 0 | 0 | 0 | 0 | 0 | 0 | 0 | 0 | 0 | |
| Ratio to Buffer | $R_P$ | $R_R$ | $-\ln R_P$ | $R_P$ | $R_R$ | $-\ln R_P$ | $R_P$ | $R_R$ | $-\ln R_P$ | $R_P$ | $R_R$ | $-\ln R_P$ | |
| | 0.025 | 1.062 | 3.689 | 0.067 | 0.833 | 2.708 | 0.017 | 0.333 | 4.094 | 0.036 ± 0.015 | 0.74 ± 0.22 | 3.5 ± 0.4 | GLU |
| | 0.027 | 1.133 | 3.624 | 0.056 | 0.722 | 2.890 | 0.014 | 0.286 | 4.248 | 0.032 ± 0.012 | 0.71 ± 0.24 | 3.6 ± 0.4 | As/Az |
| Ratio | 1.07 | 1.07 | | 0.83 | 0.87 | | 0.86 | 0.86 | | **0.92 ± 0.07** | **0.93 ± 0.07** | | |

**Table S1. Phage-specific experimental results.** Each table presents the results of experiments corresponding to a specific phage, as indicated in the upper left cell. The table is divided into four sections: three for the individual experiments and one summarizing the averages. In the upper half of the table, the first four rows correspond to either the phage of the respective table or a buffer used as a viral control, while the columns represent either susceptible or resistant bacteria to this phage, or a buffer control for the bacteria. This part of the table contains plaque counts obtained by spot titering the viral particles (PFU) in the post-cellular supernatant. The titer values are multiplied by $10^3$ PFU per milliliter (as indicated in the upper right cell), since the viruses are diluted 1000-fold during the experimental process. The results are highlighted in yellow for experiments with bacteria grown in glucose (GLU), or blue for experiments with bacteria grown in arsenate and azide (As/Az), as indicated in the rightmost column. The lower part of the table displays the ratios of viral counts from the glucose-grown (GLU) and arsenate/azide-grown (As/Az) bacterial conditions relative to the viral counts in the buffer controls. This is done for the permissive case (column $R_P$), the resistant case (column $R_R$), and the negative logarithm of the permissive case (column $-\ln R_P$), which is required for the calculation of the adsorption rate $\eta$. The final row presents the ratio of these ratios (highlighted in green), which is also depicted in Figure 2, providing insights into the differences in free virus behavior when infecting energy-competent versus energy-depleted bacterial cells for each phage. The standard error of the mean (SEM) is used to estimate the variability in the averages, accounting for the random measurement errors across the three independent experiments.

| Averages | $\lambda$ | | | $\phi$80 | | | m13 | | | T6 | | | T5 | | | x$10^3 \frac{PFU}{mL}$ |
|---|---|---|---|---|---|---|---|---|---|---|---|---|---|---|---|---|
| | Bacteria | | | Bacteria | | | Bacteria | | | Bacteria | | | Bacteria | | | |
| | Permissive | Resistant | Buffer | Permissive | Resistant | Buffer | Permissive | Resistant | Buffer | Permissive | Resistant | Buffer | Permissive | Resistant | Buffer | |
| | S3207 | S3222 | TMG | MC4100 | S2153 | TMG | S3190 | S2153 | TMG | Δxyl | tsx | TMG | MC4100 | S2153 | TMG | |
| **Virus** | 10.0 ± 0.6 | 1100 ± 208 | 1033 ± 67 | 2467 ± 467 | 11333 ± 2404 | 11333 ± 882 | 11000 ± 1000 | 30333 ± 2333 | 36000 ± 3055 | 25333 ± 3844 | 38000 ± 1528 | 42667 ± 5044 | 467 ± 176 | 10333 ± 3756 | 13333 ± 1333 | **GLU** |
| **Buffer** | 0 | 0 | 0 | 0 | 0 | 0 | 0 | 0 | 0 | 0 | 0 | 0 | 0 | 0 | 0 | |
| **Virus** | 127 ± 19 | 1267 ± 318 | 1467 ± 120 | 5900 ± 586 | 12667 ± 1856 | 10333 ± 333 | 32667 ± 333 | 35000 ± 1000 | 35333 ± 3180 | 39333 ± 1202 | 35333 ± 1764 | 41333 ± 3333 | 533 ± 240 | 11333 ± 3844 | 15667 ± 1202 | **As/Az** |
| **Buffer** | 0 | 0 | 0 | 0 | 0 | 0 | 0 | 0 | 0 | 0 | 0 | 0 | 0 | 0 | 0 | |
| **Ratio** | $R_P$ | $R_R$ | $-\ln R_P$ | $R_P$ | $R_R$ | $-\ln R_P$ | $R_P$ | $R_R$ | $-\ln R_P$ | $R_P$ | $R_R$ | $-\ln R_P$ | $R_P$ | $R_R$ | | |
| **to** | 0.0098 ± 0.0009 | 1.05 ± 0.15 | 4.64 ± 0.09 | 0.214 ± 0.024 | 0.99 ± 0.15 | 1.55 ± 0.11 | 0.307 ± 0.022 | 0.85 ± 0.06 | 1.19 ± 0.08 | 0.588 ± 0.022 | 0.91 ± 0.09 | 0.53 ± 0.04 | 0.036 ± 0.015 | 0.74 ± 0.22 | 3.5 ± 0.4 | **GLU** |
| **Buffer** | 0.087 ± 0.013 | 0.85 ± 0.18 | 2.47 ± 0.15 | 0.57 ± 0.04 | 1.22 ± 0.17 | 0.57 ± 0.07 | 0.94 ± 0.09 | 1.01 ± 0.10 | 0.07 ± 0.10 | 0.96 ± 0.06 | 0.86 ± 0.05 | 0.04 ± 0.06 | 0.032 ± 0.012 | 0.71 ± 0.24 | 3.6 ± 0.4 | **As/Az** |
| **Ratio** | **9.3 ± 2.1** | **0.81 ± 0.16** | | **2.68 ± 0.13** | **1.26 ± 0.14** | | **3.1 ± 0.4** | **1.19 ± 0.09** | | **1.64 ± 0.11** | **0.95 ± 0.05** | | **0.92 ± 0.07** | **0.93 ± 0.07** | | |
| **t(2)** | 3.95 | −1.19 | | 12.92 | 1.86 | | 5.25 | 2.11 | | 5.82 | −1.00 | | −1.14 | −1.00 | | |
| **p-value** | 0.029 | 0.36 | | 0.006 | 0.20 | | 0.034 | 0.17 | | 0.028 | 0.42 | | 0.37 | 0.42 | | |
| $\eta$ | | | **(1.55 ± 0.27) × $10^{-10}$** | | | **(1.7 ± 0.3) × $10^{-11}$** | | | **(1.32 ± 0.23) × $10^{-11}$** | | | **(3.2 ± 0.6) × $10^{-12}$** | | | **(5.8 ± 1.2) × $10^{-11}$** | |
| $\eta'/\eta$ | | | **0.53 ± 0.03** | | | **0.37 ± 0.05** | | | **0.06 ± 0.08** | | | **0.08 ± 0.11** | | | **1.03 ± 0.16** | |

**Table S2. Averages of experimental results.** This table presents the averages from the three independent experiments shown in Figure 2 and Table S1. The table is divided into five sections, one for each phage, serving as a collection of the "All Experiments Averages" section of all the respective phages from Table S1. In the upper half, the first four rows represent either the phage or a buffer used as a viral control, while the columns indicate susceptible or resistant bacteria to each phage, or a bacterial buffer control. This section presents plaque counts from spot titers of viral particles in the post-cellular supernatant. Viral titers are multiplied by $10^3$ viruses per milliliter (as noted in the upper right cell) to account for the 1000-fold dilution in the experiment. Experiments with bacteria grown in glucose (GLU) are highlighted in yellow, while blue highlights indicate growth in arsenate and azide (As/Az), as shown in the rightmost column. The lower half of the table shows the ratios of viral counts for GLU-grown and As/Az-grown bacteria relative to viral counts in the buffer controls. This calculation is performed for the permissive case (column $R_P$), the resistant case (column $R_R$), and the negative logarithm of the permissive case, which is necessary for determining the adsorption rate $\eta$. The row highlighted in green presents the ratio of these values, specifically the ratio of $R_P$ measured in arsenate and azide (As/Az) to $R_P$ measured in glucose (Glu), and similarly, the ratio of $R_R$ in As/Az to $R_R$ in Glu, with calculations structured in separate columns for clarity. This allows for a direct comparison of how the metabolic conditions influence viral counts relative to their respective controls. The standard error of the mean (SEM) is used to estimate the variability in the averages for all measurements up to the Ratio row (highlighted in green). For phage $\lambda$, this experiment corresponds to an exact replication of Brown *et al.* [2], who reported that the Ratio was greater than 1 when the host was energy-competent. Therefore, a directional hypothesis (Ratio $> 1$) was pre-specified for the energy-competent condition, and a one-tailed one-sample $t$-test versus 1 was used for this case. All other phage-host conditions were analyzed using two-tailed one-sample $t$-tests versus 1, as no prior directional expectation was assumed for those comparisons. We report the resulting $t$-values (with 2 degrees of freedom) and corresponding $p$-values below the Ratios. The last two rows, highlighted in red, present the calculated adsorption rate $\eta$, defined as $\eta = -\ln R_P/(Bt_{phage})$, where $B = (3 \pm 0.5) \times 10^9$ cells/mL represents the average number of bacteria, and $t_{phage}$ is the incubation time for each phage (with an associated uncertainty of $\pm 0.5$ min). Additionally, the relative adsorption rate $\eta'/\eta$ quantifies the effect of growth in arsenate and azide compared to growth in glucose. The uncertainty associated with the average $\eta$ and $\eta'/\eta$ are determined through error propagation, accounting for the uncertainties in the average $\ln R_P$, bacterial concentration $B$, and incubation time $t_{phage}$.

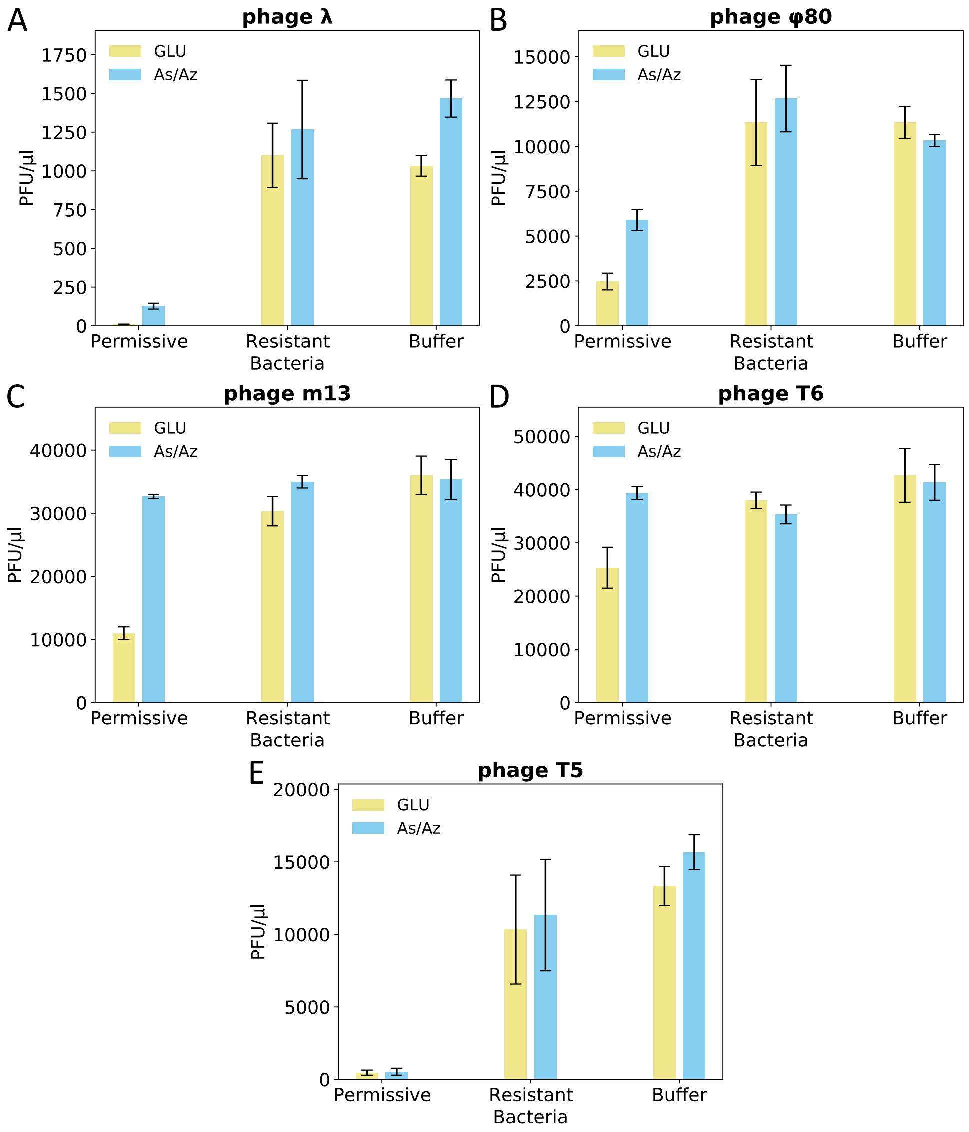


**Figure S1. Average phage counts.** Each panel displays the average count of phage particles as represented by PFU per µL in the post-cellular supernatant. Yellow bars represent phages from experiments with bacteria grown in glucose, while blue bars indicate phages from bacteria grown in arsenate and azide. Each pair of bars corresponds to interactions with different bacterial types: permissive, resistant, and buffer controls. These results are also presented in the 'All Experiments Averages' section of Table S1 and in Table S2.

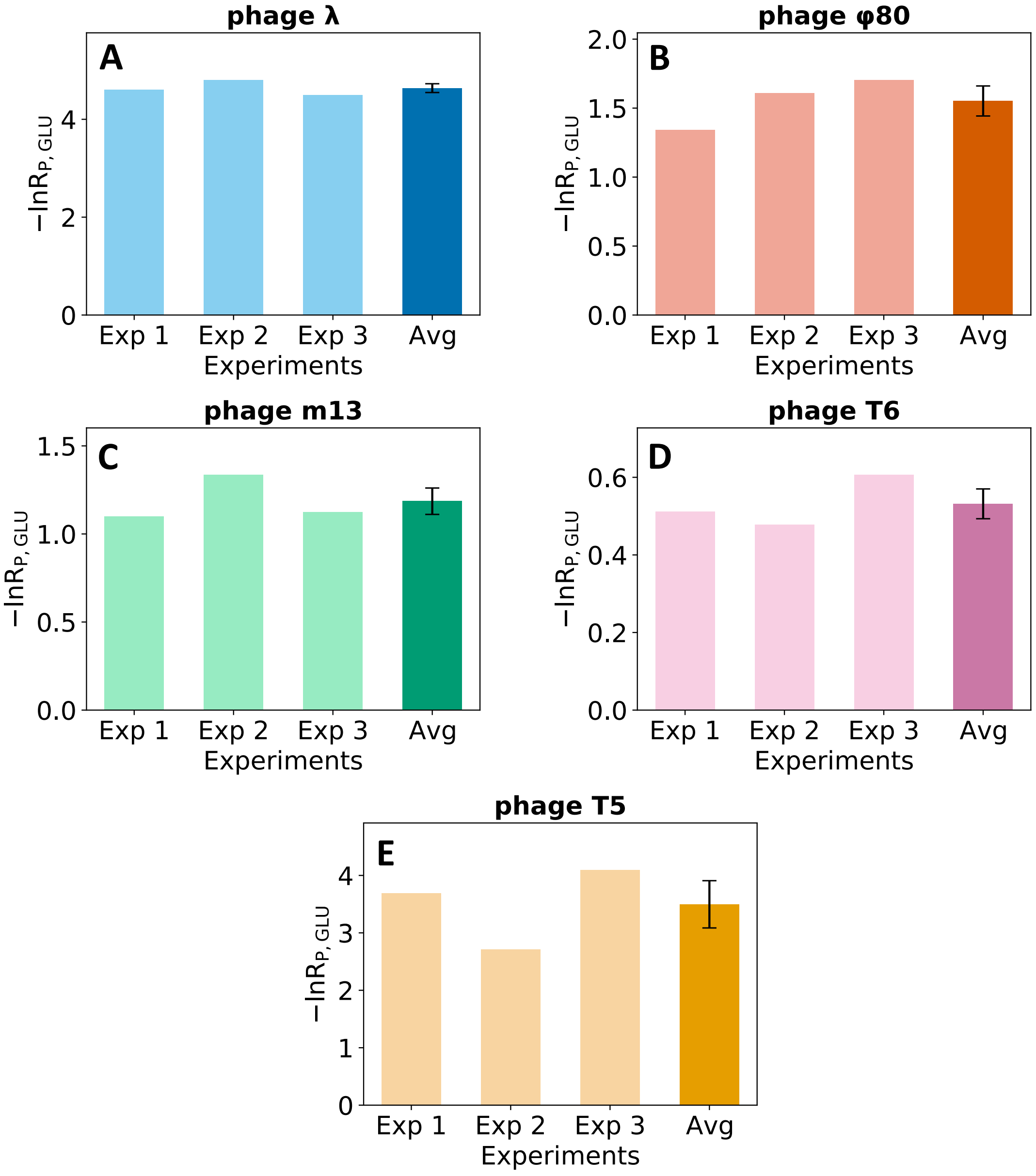


**Figure S2. Ratio to buffer in glucose across phages.** Each panel presents the results of the negative logarithm of $R_P$ in glucose, where $R_P$ represents the ratio of viral counts from permissive hosts relative to the buffer control in glucose-grown bacteria. The data are organized into four columns: one for each of the three independent experiments and one for the overall average, a necessary quantity for calculating the adsorption rate $\eta$. Each panel corresponds to a different phage, as indicated above the panel. These results are also reported in the "All Experiments Averages" section of Table S1 and in Table S2.

#### Methods and Materials

##### Strains

Bacterial strains are all derivatives of *E. coli* K12 and are listed in Table S3. Strains S2153, S3207, S3222 and S3190 are derived from MC4100 [3]. Strains tsx and Δxy are coming from the KEIO collection [1]. Phage strains are also described in Table S3 (lower part). For phage $\lambda$, we used $\lambda$wt, equivalent to $\lambda$PaPa[7], which lacks accessory tail fibers and exhibits a lower $\eta$ than the original Ur-$\lambda$ isolated by Esther Lederberg [8].

| Strain | Genotype Description |
|---|---|
| **Bacterial strains** | |
| MC4100 | F⁻ araD139 Δ(*argF-lac*)U169 flhD5301 Δ(*fruK-yeiR*)725(*fruA*25) relA1 rpsL150 rbsR22 Δ(*fimB-fimE*)632::IS1 deoC1 [3] |
| S2153 | as MC4100 but *fhuA*⁻ [10] |
| S3190 | HfrH *lacI^q^ fhuA*⁻ (this study) |
| S3207 | as S2153 but ($\lambda$*rex*::gfp) Δ(*rex-galK*)::kan [10] |
| S3222 | as S3207 but $\lambda$vir$^r$ Mal⁻ [2] |
| tsx | F⁻ Δ(*araD-araB*)567 Δ*lacZ*4787(::rrnB-3) Δ*tsx*-773::kan $\lambda$- rph-1 Δ(*rhaD-rhaB*)568 hsdR514 (name: KEIO JW0401-1) |
| $\Delta xyl$ | as KEIO JW0401-1 but tsx⁺ Δ*xylA*::kan |
| **Phage strains** | |
| $\lambda$cIb221 | $\lambda$cI ET22 Δatt b221 |
| $\phi$80vir | |
| m13 | |
| T5 | |
| T6 | |

**Table S3.** Genotypes of bacterial and phage strains used in this study. All bacterial strains are derivatives of *Escherichia coli* K12. S3222 is a spontaneous mutant resistant to $\lambda$*vir* that simultaneously lost the ability to ferment maltose, Mal⁻. tsx and $\Delta xyl$ come from the KEIO collection [1]. Phage strains include $\lambda$cIb221, $\phi$80vir, m13, T5, and T6. These phages were selected for their use of distinct entry pathways and represent a range of viral infection strategies, including lytic, lysogenic, and chronic life cycles.

##### Method and Media

Our protocol follows that of Brown et al. [2] with adjustments as specified below. We measure the adsorption of phages under two metabolic conditions: an energy-competent state (with glucose) and an energy-depleted state (with potassium arsenate and sodium azide). For each phage, the experiment is performed three times. In each experiment, bacterial cultures are initiated from independent single colonies, and phage lysates are derived from independent plaques. Exponentially growing bacterial cultures in YT broth [9] are centrifuged (1700 × g, 5 minutes, room temperature), washed with half the volume of TMG buffer (20 mM Tris-HCl pH 7.6, 5 mM $MgCl_2$, 0.1 mg/ml gelatin), and resuspended in one-fourth the volume of the same buffer.

Samples of 400 $\mu$l resuspended bacteria or TMG buffer (without bacteria) are distributed into 2-ml Eppendorf tubes. Subsequently, either 50 $\mu$l of TMG supplemented with 2% (w/v) glucose and 10 mM potassium phosphate (pH 7.4) or 50 $\mu$l of TMG supplemented with 10 mM $KH_2AsO_4$ and 0.2 M $NaN_3$ is added. The mixtures are incubated for 10 minutes at 37°C with shaking at 300 RPM. The final bacterial concentration, determined using Petroff-Hausser chamber counts, is approximately $3 \times 10^9$ cells/ml.

Phages diluted in TMG are added at MOI $\sim 0.001$ to minimize the probability of coinfections. All cultures were inoculated from the same phage dilution tube to ensure equal starting phage concentrations across conditions. After incubation with phage at 37°C and shaking at 300 RPM for a duration sufficient to allow phage adsorption while minimizing the chance of completing the first lytic cycle, bacteria are removed by centrifugation (3400 × g, 5 minutes, room temperature). The duration of incubation after phage addition was determined separately for each phage–host combination. For $\lambda$, productive lytic development was blocked in the host background used, as in Brown et al. [2]. For $\phi 80$ and T5, published latent-period values were used as guides after their compatibility with our own system had been verified [4]. M13 is a chronic filamentous phage and therefore does not have a standard lytic latent period; in our host–phage combination it required more than 1 h before phage release. For T6, we relied primarily on the kinetics observed in our own system, since adsorption was unusually slow for this phage–host pair under our assay conditions. Although literature reports describe shorter T6 latent periods under specific assay conditions [5], this is consistent with published work showing that adsorption and infection kinetics can vary substantially with host background, surface structure, and experimental conditions [6, 11]. A 100-$\mu$l portion of the supernatant is aspirated and diluted into 900 $\mu$l of TMG supplemented with 1 mM potassium phosphate (pH 7.4). Serial dilutions are prepared in TMG, and phage concentrations are determined by spotting 10-$\mu$l samples onto bacterial lawns. The fraction of unbound phage is calculated as the concentration of phage in the supernatant (PFU/ml) in bacteria-containing mixtures divided by the concentration in bacteria-free mixtures. A 100-$\mu$l portion of the supernatant is aspirated and diluted into 900 $\mu$l of TMG supplemented with 1 mM potassium phosphate (pH 7.4). Serial dilutions are prepared in TMG, and phage concentrations are determined by spotting 10-$\mu$l samples onto bacterial lawns. The fraction of unbound phage is calculated as the concentration of phage in the supernatant (PFU/ml) in bacteria-containing mixtures divided by the concentration in bacteria-free mixtures.
